# Increased EThcD efficiency on the hybrid Orbitrap Excedion Pro Mass Analyzer extends the depth in identification and sequence coverage of HLA class I immunopeptidomes

**DOI:** 10.1101/2025.06.01.657241

**Authors:** Amy L. Kessler, Kyle L. Fort, Hanno C. Resemann, Peter Krüger, Cong Wang, Heiner Koch, Jan-Peter Hauschild, Fabio Marino, Albert J.R. Heck

## Abstract

Gaining a complete and unbiased understanding of the non-tryptic peptide repertoire presented by HLA-I complexes by LC-MS/MS is indispensable for therapy design for cancer, autoimmunity and infectious diseases. A serious concern in HLA peptide analysis is that the routinely used, collision-based fragmentation methods (CID/HCD) do not always render sufficiently informative MS2 spectra, whereby gaps in the fragmentation sequence coverage prevent unambiguous assignments. EThcD can be utilized to generate complementary ion series, *i.e.* b/y ions and c/z ions, resulting in richer, more informative MS2 spectra, thereby filling in the gaps. Here, we present data generated on a novel hybrid Orbitrap mass spectrometer, facilitating fast and efficient hybrid fragmentation due to the implementation of EThcD in the ion routing multipole. We hypothesized that this would enable more comprehensive and less error-prone analysis of immunopeptidomes at minimal costs in duty-cycle. First, we optimized ETD/EThcD methods using an elastase-digested cell lysate, as this contains peptides of similar length and charge distributions to immunopeptides. Next, we compared HCD and EThcD on immunopeptidomes originating from three cell lines with distinct HLA-I complexes that present peptides with varying physicochemical properties. We demonstrate that the new instrument not only enables efficient and fast ETD reactions, but when combined with collision-based supplemental activation, *i.e.* EThcD, also consistently increases the sequence coverage and identification of peptide sequences, otherwise missed by using solely HCD. We reveal several of the biochemical properties that make HLA peptides preferably identifiable by EThcD, with internal Arg residues being one of the most dominant determinants. Finally, we demonstrate the power of EThcD for the identification and localization of HLA peptides harboring post-translational modifications, focusing here on HLA Arg mono-/di-methylation. We foresee that this new instrument with efficient EThcD capabilities enhances not only immunopeptidomics analysis, but also analysis of peptides harboring post-translational modifications and *de novo* sequencing.

## Introduction

The identification of all peptides presented by human leukocyte antigen (HLA) class I (HLA-I) complexes, collectively referred to as the HLA class I-bound ‘immunopeptidome’, is a highly valuable tool to gain insights into cellular antigen processing and presentation mechanisms under healthy and diseased conditions (1, 2). Furthermore, analysis of the immunopeptidome (i.e. *immunopeptidomics*) is crucial to uncover potential targets for the development of novel antigen-based immunotherapies (3–6). However, the low abundance, high dynamic range, and complexity of immunopeptidomes are major obstacles in the search for clinically relevant immunopeptides (7–9). Moreover, as the sought for novel antigens may originate from non-genome templated protein sequences, they can require *de novo* peptide identification strategies (10).

Currently, liquid chromatography tandem mass spectrometry (LC-MS/MS) analysis represents the most unbiased methodology for the identification of immunopeptides from either cell lines, pre-clinical models like organoids (11, 12) or primary samples such as tissues (13–17). A typical immunopeptidomics workflow includes an enrichment procedure, during which HLA-bound peptides are affinity purified, eluted from the solid phase by acid, de-salted and separated from proteins, followed by reversed-phase (RP) LC-MS analysis (18, 19). Major leaps have been made in LC-MS/MS immunopeptidomics workflows to boost identification depth through the improvement of sample preparation strategies and the introduction of different gas phase separations coupled to faster, more sensitive mass analyzers (17, 20–22). Whilst MS instruments have shown to provide sensitivity leading to in-depth analysis of immunopeptidomes, they generally don’t allow for fragmentation flexibility covering the full physicochemical space (23–25).

LC-MS/MS-based identification of immunopeptides is currently predominantly accomplished by MS sequencing using solely collision-based techniques, such as collision induced dissociation (CID) or higher-energy collision induced dissociation (HCD) (26, 27), which provide the advantages of high fragmentation speed, and optimized database search strategies. Both CID/HCD have demonstrated excellent effectiveness for identifying tryptic peptides generated from proteome digestion, resulting in high identification rates (28). Immunopeptidomes, however, are complex mixtures of non-tryptic peptides, which are typically 8-12 amino acid residues in length and elute from up to six different HLA alleles. Within their binding groove, HLA-alleles favor the presence of certain amino acids at position two (P2) and the C-terminus (C-term) of the peptide sequence, termed the anchor residues, and to a lesser extent at other positions, termed sub-anchor residues (29). This amino acid preference of anchor and sub-anchor residues makes up the so-called *binding motif* of each specific allele. The presence of allele-specific binding motifs, i.e. different amino acid sequences, adds an additional layer of complexity to immunopeptidomics analysis and potentially yielding differences in fragmentation efficiency across the varying motifs. Therefore, depending on the fragmentation scheme used, only a limited portion of the obtained MS2 spectra might provide adequate sequence-diagnostic information in immunopeptidomics. This may impact the accuracy of peptide sequence assignments through database searches, post-translational modifications (PTMs) identifications (30), and further hinder the accuracy of *de novo* search methodologies (31).

A hybrid fragmentation scheme termed “electron-transfer/higher-energy collision dissociation” (EThcD) has been previously introduced (32). This method isolates a single ion package and employs dual fragmentation to generate both the fragment ion series induced first by ETD (c/z) and then by HCD (b/y) in a single MS2 spectrum. The generation of dual-fragment ion series results in more complete MS2 spectra that facilitate highly confident peptide assignment and localization of PTMs (33–35). We have previously shown the advantage of EThcD for the analysis of immunopeptides and their associated PTMs (36–39). Nevertheless, the widespread implementation of EThcD has stayed behind in the field of immunopeptidomics, likely due to the requirement of costly instrumentation and the need of complex method settings needed for the optimized fragmentation of samples with widely diverse immunopeptidomics features. In addition, fragmentation completeness and reaction speed in ETD are highly dependent on the charge density of peptide precursors (40). Importantly, due to their relatively small size and non-tryptic cleavage, immunopeptides mainly exhibit charge states (*z*) between +1 and +3 in gas phase (26), posing a limitation to EThcD fragmentation completeness and requiring long reaction times, ultimately impacting the overall duty cycle (41, 42).

While the Orbitrap™ Exploris™ mass spectrometer has been well received, its fragmentation capabilities have so far been limited to HCD. Here, we describe the use of a novel hybrid Orbitrap-based mass analyzer, termed the Orbitrap™ Excedion™ Pro, which features additional ETD and EThcD capabilities. In this instrument, EThcD and ETD can be performed in the ion routing multipole (IRM), exploiting its substantial length for high ion storage capacity. With these higher ion densities, the ETD reaction times can be shortened even for peptides with lower charge state, leading to reduced duty cycles while profiting of the complementary sequence coverage provided by EThcD fragmentation.

To explore the capacities of the new instrument for immunopeptidomics, we first fine-tuned MS methods inclusive of the new ETD and EThcD ability. To this end, we used an elastase non-tryptic digest which closely resembles the global biophysical features of an HLA-I immunopeptidome. We then analyzed a set of HLA-I immunopeptidomes purified from three cell lines displaying peptides presented by a variety of different HLA-I alleles, thus resulting in physicochemically distinct peptide ligandomes, using EThcD and HCD. We demonstrate that by using EThcD on this new instrument, higher peptide sequence coverage can be obtained compared to HCD across a wide range of HLA-I alleles, and that it uncovers a part of the immunopeptidome that HCD fails to identify. We report several peptide features where EThcD outperforms HCD in terms of sequence identification, with internal Arg residues being an important discriminator. Lastly, we complemented our analysis by demonstrating that a set of peptides harboring mono-/di-methylation are also impacted by the “blindness” of HCD for specific peptide sequences, and that these are more readily identified by EThcD.

## Experimental procedures

### Experimental design and statistical rationale

A description of the experimental design including sample naming, an overview of run conditions, RAW MS file names and the number of biological/technical replicates can be found in **Supplemental Tables 1-3**, as well as the raw Fragpipe psm output. The acquired data for this study was generated as follows: one batch of HeLa elastase digest was used for ETD method optimization and for each cell line one biological replicate was harvested and subjected to HLA-I immunopurification. The elastase digest and enriched immunopeptides were divided in such manner to have two technical replicates per measuring condition to ensure reproducibility. Motif analyses of fragmentation-preferred amino acids were performed in Two-Sample Logo using the binomial test (twosamplelogo.org; (43, 44)). Residues are shown if p-value<0.05.

### HeLa digest preparation and enrichment of HLA class I peptides

A HeLa cell pellet was lysed with 8M urea in 100mM Tris. The lysate was prepared by incubation with 5mM dithiothreitol (DTT) at 56°C for 25 min followed by incubation with 20mM iodoacetamide (IAA) for 45min in the dark at room temperature. Prepared lysates in 2M urea in 100mM Tris were then digested with 2μg/mL elastase for 4h at 37°C. Digestion was quenched with a final concentration of 0.5% TFA before C18 clean-up. JY, GR and Jurkat cells were cultured in RPMI-1640 (Capricorn) supplemented with 10% fetal calf serum (Gibco) and 1% penicillin-streptomycin (100 U/mL, 100μg/mL; Gibco) at 37°C, 5% CO2. Dry cell pellets were stored at -20°C until further processing. HLA-I-bound peptides were enriched as previously described (45). In short, cell lysates were incubated with anti-pan-HLA-I (mAb clone W6/32 (Bio-Legend) coupled toProtein A sepharose Fast Flow (Cytiva) affinity beads for 3h at 4°C. HLA-I-bound peptides were eluted with 10% (v/v) acetic acid followed by C18 desalting. Peptides were eluted from the C18 cartridge (Waters) in two separate fractions of 25% and 80% ACN respectively. The 80% ACN peptide fraction was further cleaned up by 5kDa MWCO filtration (Merck/Millipore). Enriched peptides were combined, dried down and resuspended in 1% ACN, 0.1% TFA prior to LC-MS/MS injection.

### The Orbitrap Excedion Pro mass spectrometer

A schematic overview of the new mass spectrometer is shown in **Fig. 1**. Electron-transfer dissociation (ETD) with the option for supplemental collisional activation, i.e. electron-transfer/higher-energy collision dissociation (EThcD), can be performed inside the IRM of the instrument. To this end, after precursor ions have been injected into the IRM, reagent anions are created using the EASY-ETD™ ion source and transferred into the same trapping cell. Reagent and precursor ions are separated by means of an axial DC gradient along the IRM. To start the ETD reaction, this gradient is then switched off and RF voltages are applied to the front and exit electrode of the IRM to trap reagent anions and precursor cations simultaneously. After the desired reaction time, the resulting precursor and fragment ions are transferred to the C-Trap. Subsequentially, the ion package can be directly injected into the Orbitrap mass analyzer for mass analysis (ETD-mode) or be collisionally activated by acceleration into the IRM (EThcD-mode). Optimal reaction times are automatically calculated and set based on the precursor charge state (41).

**Figure 1.**
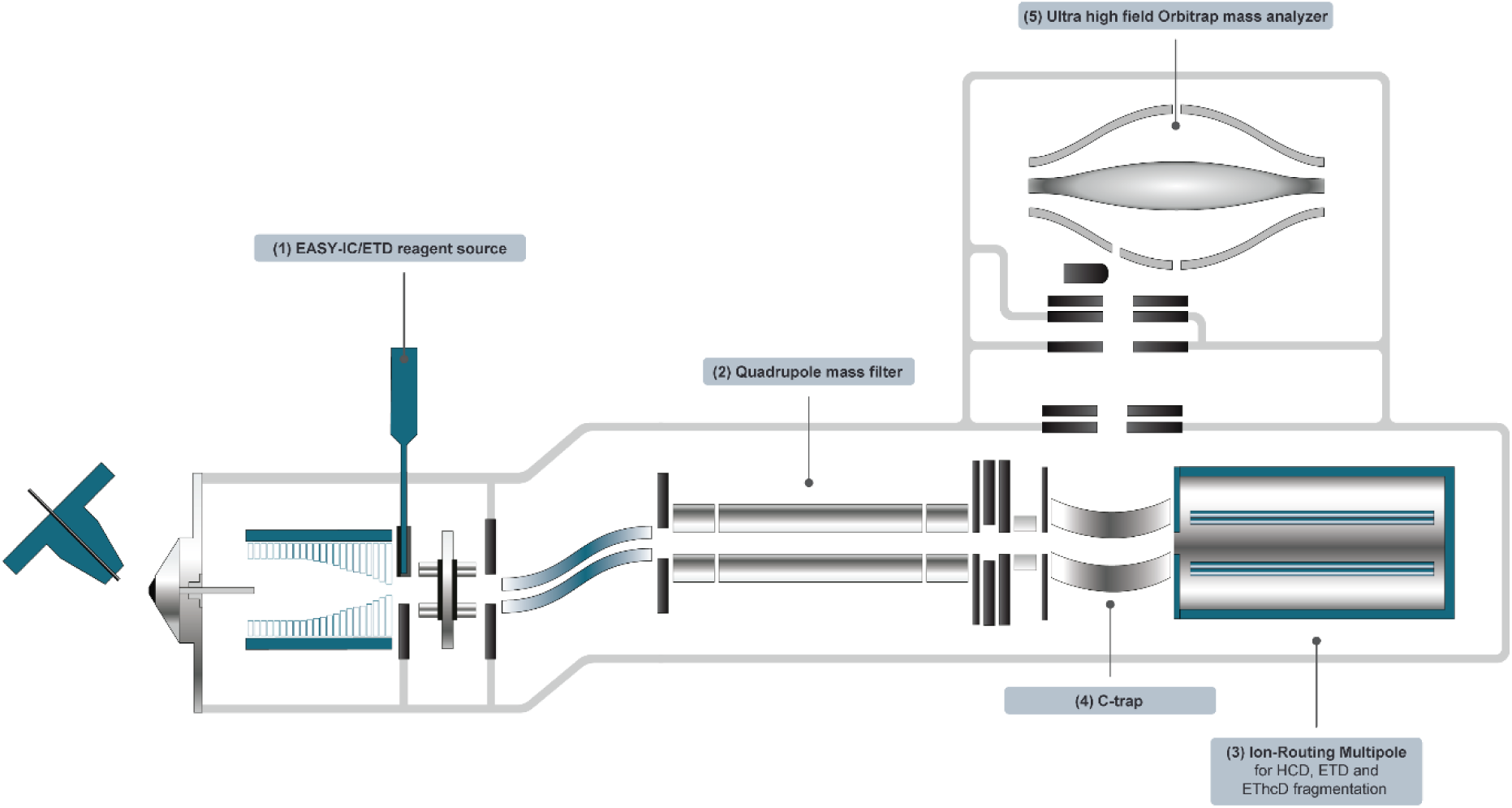
Schematic of the Orbitrap Excedion Pro mass spectrometer with ETD and EThcD capabilities. In particular, the new features of the mass spectrometer are highlighted, including the (1) EASY-IC/ETD™ reagent source enabling Electron-transfer dissociation (ETD), (2) quadrupole mass filter for selecting ion packets and sending for fragmentation, either HCD, ETD or EThcD, in the (3) Ion-Routing Multipole. The resulting fragments are then passed to the (4) C-Trap before injection in the (5) ultra high field Orbitrap mass analyzer.

### Liquid chromatography-mass spectrometry (LC-MS/MS) analysis

All samples were analyzed by nanLC-MS/MS using a Thermo Scientific™ Vanquish™ Neo and an Orbitrap Excedion Pro mass spectrometer using research-grade instrument control software. A one-column set-up was used with a 25cm x 0.0075cm inner diameter analytical column with integrated emitter (IonOpticks). For the elastase digest injections, peptides were separated on a 24 min acetonitrile in water gradient (4-40%) at a flow rate of 300 nL/min. Immunopeptidomes were separated on a 56 min acetonitrile in water gradient (4-45%) at a flow rate of 300 nL/min. The solvents contained 0.1% FA in water and 80% acetonitrile. Full MS and MS2 spectra were acquired in an Orbitrap mass analyzer at a resolution of 60,000 and 15,000, respectively. The variable parameters, including *m/z* range, supplemental activation (SA; for EThcD), normalized collision energy (NCE; for HCD), precursor charge inclusion for MS2, and RF%, used for the elastase digest and immunopeptidome analysis are listed in **Supplemental Table 1 and 2**. All samples were measured using calibrated charge-dependent ETD reaction times, aiming for a precursor depletion of 13.5 % of the precursor abundance (41). AGC was set to 50% for for both HCD and EThcD experiments and the maximum injection time at 120ms. For the method optimization, 60ng elastase digest was injected. For the immunopeptidomics analyses, immunopeptides derived from the equivalent of 3.5e7 JY cells or 1.8e7 GR/Jurkat cells were injected.

### Data analysis

Analysis of MS data was performed by using Fragpipe v22.0. Data were searched against the human proteome (UP000005640, containing 20590 number of protein sequences) with the addition of 50% decoys in the search space and non-specific enzymatic digestion. Precursor ion and MS2 tolerances were set to 20 ppm. For the elastase digest analysis, peptides between 7-25 amino acids long and a mass between 700-3000 were considered. Default variable and fixed modifications within the non-specific HLA workflow were included, with the addition of the fixed modification of carbamidomethylation on cysteine. For the analysis of the immunopeptidomes acquired in this study, peptides between 8-15 amino acids long and a mass between 500-3000 were considered. The following variable modifications were allowed: methionine oxidation, N-terminal acetylation, N-terminal pyro-glu, N-terminal glutamate to pyroglutamate conversion, cysteinylation. For the mono- and di-methylation on arginines, a separate analysis was performed while utilizing the same parameters as stated above for the immunopeptidomes, with the exception of a selected mass range between 700-1500. Fixed modifications default to the workflow were included and localization score thresholds were calculated by PTMProphet (46). For the analysis of all public data, peptides between 8-15 amino acids long and a mass between 700-1500 were considered. Default variable and fixed modifications within the non-specific HLA workflow were included. All results were filtered to <1% FDR at the peptide level for stringent cut-off.

HLA binding predictions were performed with NetMHCpan 4.1 (47), considering peptides as binders with a percentage elution rank below 2% unless differently stated for specific analyses. Sequence motifs of allele-assigned peptides were plotted using WebLogo (https://weblogo.berkeley.edu/logo.cgi; (43, 44)). For further bias analyses and comparisons between EThcD and HCD of immunopeptidomics data across the 3 cell lines, all the identified peptides were filtered for more stringent elution ranking (<1%) to increase the allele specificity assignment and decrease confounding factors. Peptide overlaps between EThcD and HCD, calculation and display of chemical physical properties such as charge state and length distributions were performed in GraphPad PRISM v.10. To study sequence biases between HCD and EThcD, motif analysis of fragmentation-preferred amino acids per cell line and HLA-allele were performed in Two-Sample Logo using the binomial test (twosamplelogo.org; (43, 44)). Residues are shown if p-value<0.05.

Extraction and assignment of ion series and sequence coverage from HCD and EThcD were performed with an in-house written Python script. In short for sequence coverage calculations, in-silico theoretical fragment ion lists were created for each detected precursor primary sequence and charge state as denoted from the PSM outputs of FragPipe v22.0 using a custom script utilizing the Pyteomics python library (48, 49). For HCD-fragmentation, b and y ion types were considered. For EThcD-fragmentation, b, y, c and z-dot ion types were considered. Each PSM was then initially searched against the respective theoretical fragment list and matching signals were determined using a 10 ppm mass tolerance and a match between the theoretical charge state and the charge state annotation by the acquisition software. If no matching fragments were found using this strict criterion, a second search was conducted whereby the requirement of charge state annotation was removed, but the PPM filter requirement was increased to 5 ppm. The second search is used to minimize false negatives as charge state annotation by the acquisition software may sometimes be absent, particularly for peaks with low signal-to-noise (SNR). If neither of these searches identified any matched fragments, these precursors were excluded from further processing.

## Results

### Using an elastase cell digest as an immunopeptidome surrogate for EThcD method optimization on the new instrument

EThcD has previously been shown to extend the detectability of immunopeptidomes (39). However, its use has lagged behind in the immunopeptidomics field due to the slower duty cycle compared to HCD as well as the lower fragmentation efficiency for immunopeptides exhibiting low charge density (42). As one of its new features, EThcD in the Orbitrap Excedion Pro mass analyzer is now performed inside the IRM (**Fig 1**). In this way, high ion densities can be utilized, and short cycle times for low charge state precursors are facilitated. Specifically, reaction times as short as 13 ms and 6 ms were utilized for precursor charge states z=+2 and z=+3, respectively. To evaluate its potential and optimize the new fragmentation feature of the instrument, we initially analyzed an elastase cellular digest, as surrogate for an HLA ligandome sample, which is significantly more precious. Elastase is an easy-to-use, low-specificity digestion enzyme which, upon fine-tuning of the digestion conditions, can yield peptides of similar charge state (*z*; generally 1-3), *m/z* distributions and length (typically 8-15 amino acids) to the ones binding to HLA-I complexes (26, 50). With the elastase digest, we optimized supplemental activation (SA) in EThcD, charge state (*z*) selection for MS2, *m/z* scan ranges, and source transmission radio frequency (RF) (**Supplemental Table 1**). For the evaluation, we used the Hyperscore within FragPipe as a metric for identification confidence. We selected method parameters tuned towards similar length and charge distribution to immunopeptidome analysis.

First, ETD experiments were performed without any supplemental activation. Next, given the importance of sequence coverage for immunopeptidomics and the complementary nature of EThcD fragment ions, we performed EThcD at various SA, ranging from 15 to 35% (**Fig. 2A**). This SA range was selected based on previous reports where the optimum was found to be SA30% for immunopeptidomic analyses (51). Compared to ETD-only (i.e. no SA activation) fragmentation, EThcD fragmentation increased the total number of identified PSMs, with the highest PSM identifications observed at SA25% and 30%. When compared to ETD only, the gain in PSM numbers with SA≥25% can be explained by the additional identification of *z*=+2 peptides, which without SA mainly undergo non-dissociative electron transfer rather than backbone fragmentation (40). SAs 25% and 30% behaved very similarly and achieved the highest number of PSMs and Hyperscore assigned by Fragpipe. Given the previous use of SA30% for immunopeptidomics (36, 37, 50), and nearly no observed differences between SA25% and SA30%, we selected SA30% for the subsequent immunopeptidome measurements.

**Figure 2.**
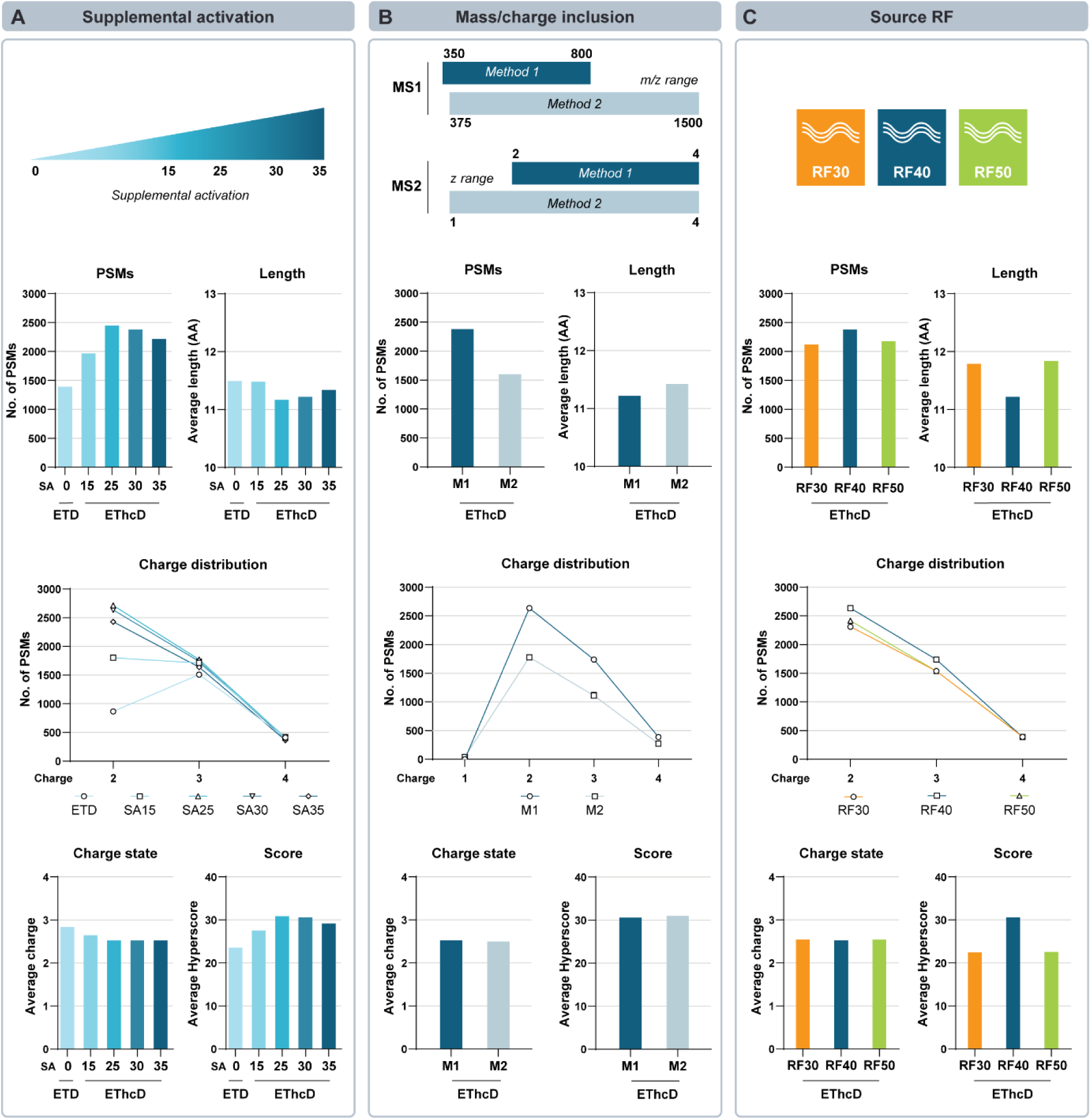
Optimization of EThcD methods on the new Orbitrap Excedion Pro using an elastase HeLa cell digest. An elastase HeLa cell digest was used as this generates peptides with features similar to those present in an immunopeptidome. The metrics used for the optimization included number of identified PSMs, peptide length and charge distributions, average peptide charge state and identification (ID) confidence scores (Hyperscore). At the top is schematically depicted which experimental parameters were varied: supplemental activation (SA), precursor ion mass and charge inclusion, and percentage of RF amplitude applied to ion funnel (RF). **(A)** Optimal SA was determined by comparing ETD-only and EThcD with increasing degrees of SA: 15%, 25%, 30% and 35%. **(B)** Two methods (M1 and M2) with different *m/z* and *z* ranges were tested. For M1 an *m/z* range between 350-800 and *z* range of 2-4 was used, whereas for M2 an *m/z* range between 375-1500 and *z* range of 1-4 was used. **(C)** Three RFs were tested at 30%, 40% and 50% for optimal precursor distribution.

Next, methods were optimized on the elastase sample for the inclusion range of *m/z* and *z* values (**Fig. 2B**). Method 1 (M1) consisted of a narrower *m/z* range 350-800 and included *z*=+2-4. While several reports have shown that HLA-I peptides, depending on the alleles expressed, can display a noticeable percentage of *z*=+1 peptides, we reasoned that a narrower *m/z* range and exclusion of *z*=+1, which could otherwise include the potential fragmentation contaminants, would increase method sensitivity and increase identification rates. Method 2 (M2), on the other hand, permitted scanning in the wider range between 375-1500 *m/z* and included singly charged precursor selection for MS2. M1 yielded higher PSM numbers than M2 (+49%). Additionally, there is almost no detectable differences with regards to the peptide average length, charge state distribution or Hyperscore (**Fig. 2B**). Interestingly, despite the inclusion of singly charged precursors, very few *z*=+1 peptides were identified with M2. We reasoned, however, that immunopeptidomes can still contain lower *z* peptides than the ones present in our elastase digests. For that reason, we decided to continue with both M1 and M2 methods for subsequent JY immunopeptidome measurements, as it had been previously reported that JY immunopeptidomes can contain up to 35% singly charged peptides, albeit that this observation was made in a different mass spectrometry platform (52).

Lastly, three different source conditions were compared by varying the amplitude of the RF voltage applied to the ion funnel (**Fig. 2C**). Except for length and Hyperscore assignments, only marginal differences were observed in the elastase digest identifications between all three used RF values. Considering the length of HLA peptides will mostly be clustered around 9-mers, and higher Hyperscores indicate higher confident identifications, RF40% was selected for immunopeptidome measurements. Taken together, the elastase digest served as a valuable surrogate immunopeptidome sample that allowed for method optimization, and based on this data SA30%, M1 and RF40% were optimized and selected for subsequent analyses of the HLA ligandomes.

### Fragmentation propensities are demarcated by the physicochemical properties of HLA allele-specific immunopeptidomes

Considering the HLA allele-specific physicochemical properties of immunopeptides, we hypothesized that any blind spots by HCD, and thus the complementary value of EThcD, could be sequence-specific and therefore HLA allele-driven. To understand such potential fragmentation biases, we first interrogated the physicochemical characteristics of HLA-bound peptides in a set of publicly available immunopeptidomics data acquired by HCD, containing matched HLA profiles to the cell lines used in our study (**Supplemental Table 3**) (14, 50, 53, 54). Once the data was searched with FragPipe, the HLA specificity of peptides was predicted by NetMHCpan 4.1 (47) and only predicted binders with a rank <2% were further considered for the HLA-A*01:01, -A*02:01, -A*03:01, - B*07:02, -B*27:05 and -B*35:03 alleles. HLA-C alleles were omitted from this analysis due to their relatively small contribution to the overall presented immunopeptidomes and the potential promiscuity of their binding motifs (55).

As ETD is known to have higher fragmentation efficiency for peptides with a higher charge and charge density, we first looked at the length, average charge state and calculated the charge density for the allele-specific immunopeptidomes. In line with previous reports (56, 57), HLA-A*01:01, -A*03:01 and -B*27:05 alleles allow longer peptides within their binding pockets (**Fig. 3A**). Of those, HLA-A*03:01 and -B*27:05 harbor the highest proportion of *z*=+3 peptides (31% and 42%, respectively), whereas HLA-A*01:01 and -B*35:03 harbor the highest percentage of *z*=+1 peptides (27% and 40%, respectively) (**Fig. 3B**). From these length distribution and charge states, the average charge density was calculated (average charge/average no. of amino acids). Indeed, HLA-A*03:01 and -B*27:05 contain peptides with the highest charge density, potentially facilitating immunopeptides identification by EThcD fragmentation (**Fig. 3C**).

**Figure 3.**
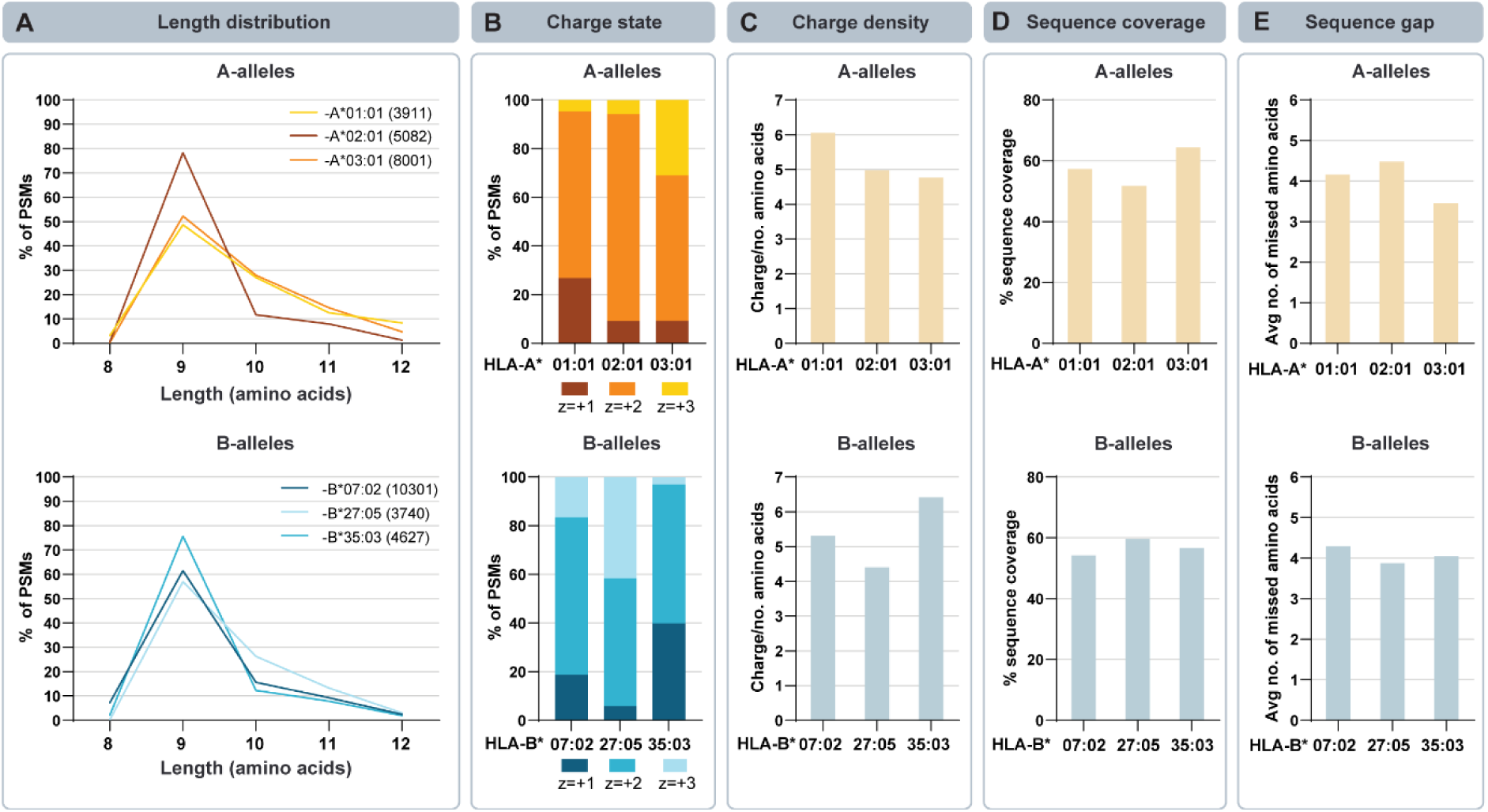
Immunopeptidome features, as extracted from publicly deposited HCD-only datasets, are HLA allele-specific. We selected datasets from deposited data (**Supplemental Table 3**) that were matched to the HLA alleles present in the JY, GR and Jurkat cell lines investigated in this study. Analysis was performed on the HCD-only identified peptides that were predicted binders (<2% binding rank by NetMHCpan 4.1) to the HLA-alleles. For analysis, the following HLA alleles were included (PSM numbers in brackets): HLA-A*01:01 (3911), -A*02:01 (5082), -A*03:01 (8001), -B*07:02 (10301), - B*27:05 (3740), and -B*35:03 (4627). Various features such as **(A)** peptide length distributions (8-15 amino acids) and total number of peptide entries used, **(B)** average charge states, **(C)** average charge density (defined as the average ratio of gas phase-detected charge to peptide length), **(D)** average sequence coverage and **(E)** average sequence gap (defined as the average number of missed amino acids) were extracted from the deposited data and calculated to uncover HLA allele-specific features driving HCD fragmentation biases.

To uncover potential HCD biases driven by the presence of certain amino acids in immunopeptides, we calculated the average sequence coverage for each allele-specific immunopeptidome. The average sequence coverage reached by HCD was highest for A*03:01 and B*27:05 (**Fig. 3D**). This likely is facilitated by the resemblance of their immunopeptides to tryptic peptides with a charge on both the N- and C-terminal residues (58). As another indicator of sequence ambiguity assignment, we calculated the sequence gap, which is defined as the average number of amino acids not covered by any ion fragment (**Fig. 3E**). HCD yielded a maximum average sequence coverage of 64% for the tryptic-like HLA-A*03:01 peptides and a minimum average sequence coverage of 52% for -A*02:01 peptides. Similarly, the sequence gap of HLA-A*02:01 peptides is 4.5 amino acids, leaving a high level of ambiguity for this set of highly clinically relevant peptides (59–62).

### EThcD decreases sequence ambiguity and uncovers part of immunopeptidome missed by HCD

We next sought to apply our optimized methods on the new mass analyzer with expanded ETD capabilities, and interrogated HLA class I ligandomes from three distinct cell lines covering a wide range of HLA alleles (**Supplemental Table 2**). These cell lines were not only selected because of their potential clinical relevance, as they collectively cover most of the world population in terms of HLA allele expression (63–65), but also to further understand how EThcD can help uncover the “HCD-undetectable” immunopeptidome across a wide range of HLA alleles (**Table 1**). The JY cell line, a B cell lymphoblastoid line, is homozygous for HLA-A*02:01, -B*07:02 and -C*07:02, and is frequently used in immunopeptidomics research (15, 66). Immunopeptides were enriched from this cell line and analyzed by LC-MS/MS on the new instrument. The length distribution analysis revealed a high enrichment for 9-mers, which is the preferred length for HLA-binding peptides (**Supplemental Fig. S1A**). Furthermore, over 90% of the isolate was predicted by NetMHCpan 4.1 to bind at least one HLA the cell line’s HLAs below the commonly used binding threshold of 2%, further corroborating that we had a pure sample of bona fide HLA binders (**Supplemental Fig. S1B**). Starting with this high-quality JY immunopeptidome, we first evaluated differences between method M1 and M2 (**Fig. 2B**) for both HCD and EThcD fragmentation.

**Table 1.** Overview of studied cell lines with the respective 4-digit human leukocyte antigen (HLA) class I alleles they present.

| Cell lines | HLA-A* | HLA-A* | HLA-B* | HLA-B* | HLA-C* | HLA-C* |
| --- | --- | --- | --- | --- | --- | --- |
| JY | 02:01 | 02:01 | 07:02 | 07:02 | 07:02 | 07:02 |
| GR | 01:01 | 03:01 | 07:02 | 27:05 | 02:02 | 07:02 |
| Jurkat | 03:01 | 03:01 | 07:02 | 35:03 | 04:01 | 07:02 |

EThcD fragmentation, independent of the method used, yielded a higher average Hyperscore for peptide identification when compared to HCD, again likely due to the effect of complementary b/y and c/z ion series in EThcD (+21% and +15% higher average Hyperscores for M1 and M2, respectively) (**Fig. 4A**). Independently from the method used, a pronounced difference between EThcD and HCD was observed for the reached ID rate, defined as the percentage of identified PSMs from the total number of MS2 scans (a total of +16.3% and +11.3% for M1 and M2, respectively) (**Fig. 4B**). The higher proportion of MS2 scans assigned to PSMs of EThcD compared to HCD fragmentation alone suggested that the dual b/y and c/z ion series benefits spectral quality, density and subsequent “gap-free” sequence assignment. As also shown by the elastase data, M1 rendered a higher proportion of identified PSMs and average score with both EThcD and HCD, even in the JY cell line where more *z*=+1 peptides are expected (15% *z*=+1 in our dataset). Worth of note, of all *z*=+1 peptides in our dataset, 75% of the precursors were also found in higher charge states and still included for analysis with M1. Thus, we selected M1 for immunopeptidome analyses of the other two cell lines.

**Figure 4.**
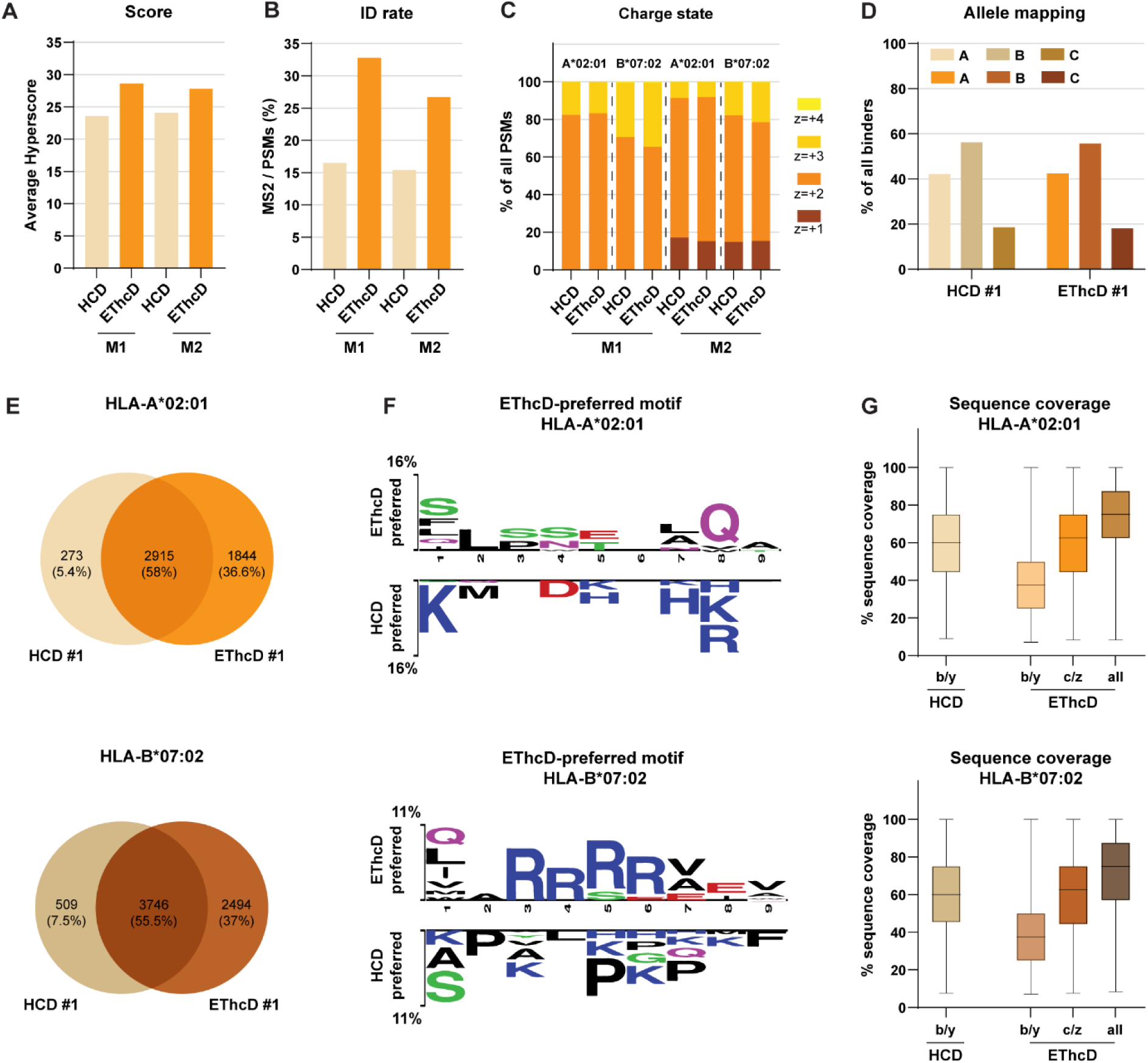
EThcD fragmentation increases sequence coverage and enables identification of JY immunopeptides missed by HCD. For JY immunopeptidome analysis, methods M1 and M2 were compared with both HCD and EThcD for **(A)** the resulting spectral score (Hyperscore) and **(B)** ID rates. Identified predicted binders (NetMHCpan 4.1, <1% binding rank) were extracted for HLA-A*02:01 and –B*07:02 and analyzed for their **(C)** charge state distribution and **(D)** percentage (%) of binders to the HLA-A, -B, and –C-alleles. N.B. Identified predicted binders can be predicted to bind multiple HLA-alleles. **(E)** Venn diagrams of the overlap between HCD- and EThcD-identified unique peptides for HLA-A*02:01 and –B*07:02. **(F)** Sequence motifs of over-represented amino acids in identified peptides. Motifs on the top are enriched in EThcD-preferred peptides, whereas motifs on the bottom are less identifiable in the EThcD-preferred peptides compared to sequences identified by HCD alone + those identified by both HCD and EThcD (i.e. ‘HCD-preferred’). Motifs were extracted with Two-Sample Logo (p<0.05) for HLA-A*02:01 and –B*07:02 predicted binders (<1% binding rank). The height of each symbol is proportional to the difference in relative frequencies. **(G)** Percentage (%) of sequence coverage of HCD (b/y ion series) and EThcD (b/y, c/z, or the combination of ion series) for HLA-A*02:01 and –B*07:02 predicted binders (<1% binding rank).

We next set out to delineate from which alleles and to what extend EThcD fragmentation can identify complementary peptide sequences. Irrespective of fragmentation (HCD vs. EThcD) and method (M1 vs. M2), all JY peptides were identified to be predominantly doubly charged although HLA-B*07:02 peptides had a higher proportion of triply charged peptides than HLA-A*02:01 (**Fig. 4C**). Identified peptides were mapped back to each host cell HLA allele based on NetMHCpan 4.1 binding predictions (**Fig. 4D**). To prevent motif ambiguity, only peptides below the stringent binding threshold of <1% were considered for subsequent analysis. Most peptides were derived from the B-alleles, followed by the A- and C-alleles, respectively. No differences were observed in allele-designation between HCD and EThcD, nor were there differences in the binding motifs of the peptides (**Supplemental Fig. S2**).

Out of the total 5032 unique HLA-A*02:01 stringent binders identified in JY cells, 1844 (37%) were EThcD-specific and 273 (5%) were HCD-specific in this dataset (**Fig. 4E**). Motif enrichment analysis (43, 44) of the EThcD-specific peptides compared to the HCD-specific and overlapping peptides revealed a subtle enrichment of hydrophobic, aliphatic amino acids (mainly Leucine (Leu; L) and Serine (Ser; S)) in the “EThcD-preferred” sequences (**Fig. 4F**), while HCD-identified peptides displayed a clear bias for charged amino acids close to the C- and N-termini (i.e., tryptic-like). Importantly, EThcD dual fragmentation boosted sequence coverage in the HLA-A02:01 dataset to an median of 75% when compared to the 60% in HCD (**Fig. 4G**).The same analysis was performed for HLA-B*07:02-predicted peptides, of which 2494 (37% of the total 6749) were EThcD-specific in our dataset, with just 509 (7.5%) being specific for HCD (**Fig. 4E**). Here, motif enrichment analysis unveiled a clear dominance of primarily the basic amino acid arginine (Arg; R) somewhere close to the middle of the peptide sequence. Notably, this data aligns well with previous work that showed that such internal Arg residues hamper CID/HCD-fragmentation due to a general decrease in proton mobility and a consequently limited yield of fragments across the peptide backbone (**Fig. 4F**) (67). Instead, prolines (Pro; P) seem to be underrepresented in the EThcD dataset, likely due to the effect of Pro causing low incidence of ETD dissociations (68). Notably and similarly to HLA-A*02:01 peptides, the median peptide sequence coverage for EThcD-identified peptides was substantially higher (75%) than for HCD-identified peptides (60%) (**Fig. 4G**).

To corroborate our findings that EThcD expedites unique identifications, we referenced our data against a very large set of publicly available HCD datasets of immunopeptidomes of JY cells (cumulatively 114 runs consisting of 19849 distinct peptide sequences; **Supplemental Table 3**; (52, 66, 69)) and cross-referenced all these HCD-identified peptides to our current list of 1844 HLA-A*02:01 and 2494 HLA-B*07:02 EThcD-preferred peptides. Using this comprehensive dataset we uncovered the peptide sequences that were exclusively detected by EThcD fragmentation and would be missed with the sole use of CID/HCD fragmentation measurements, even after 114 independent runs, i.e., the reference dataset used here (**Supplemental Fig. S3A**). Motif enrichment analysis for the HLA-A*02:01- (**Supplemental Fig. S3B**) and -B*07:02-mapped EThcD-preferred peptides (**Supplemental Fig. S3C**) again demonstrated a strong presence of Arg residues near the center of the peptide sequence, when compared to peptides that were identified solely by using HCD. Additionally, there was an enrichment of Leu in P5-P8. Taken together, our data indicate that EThcD is highly beneficial for the identification of centric Arg-containing HLA peptides. As such peptides are over-presented in specific alleles, for instance in HLA-B*07:02-derived immunopeptidomes, and to a lesser extend in -A*02:01-derived immunopeptidomes, EThcD is essential to cover allele-specific immunopeptidomes more comprehensively.

### EThcD fragmentation extends the detectable immunopeptidomes for a wide range of HLA alleles

Having demonstrated the added value of EThcD fragmentation for JY immunopeptidome identification, we next aimed to extend our observations to a wider range of alleles. To this end, we selected the GR (B-lymphoblastoid) and the Jurkat (T cell leukemia) cell lines (**Table 1; Supplemental Table 2**). For these samples, quality control analyses such as length distribution analysis and HLA binding prediction, also demonstrated that our experiments provided pure immunopeptidome samples of bona fide HLA binders (**Supplemental Fig. S1**).

As observed for the JY immunopeptidome, EThcD fragmentation spectra from the GR immunopeptidome displayed higher identification confidence (+25% higher Hyperscore) (**Fig. 5A**), and identification rates when utilizing EThcD fragmentation instead of HCD (a total of +16.4%) (**Fig. 5B**). Using the stringent prediction threshold of <1% rank by NetMHCpan 4.1, peptides were mapped to each donor HLA allele (**Fig. 5C**) and again the sequence motifs of allele-specific binders were plotted (**Supplemental Fig. S2**).

**Figure 5.**
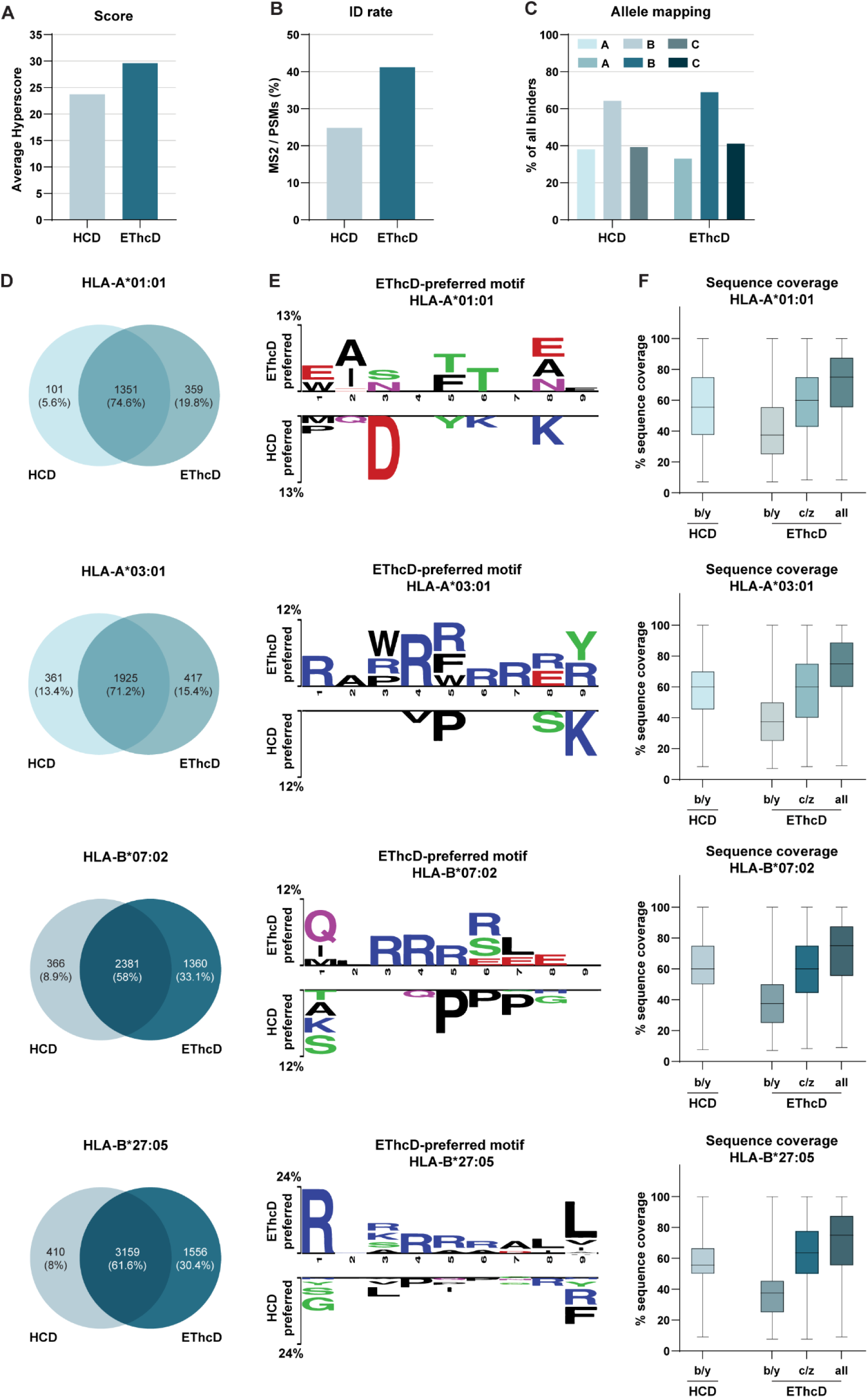
EThcD fragmentation extends the detectable immunopeptidome of the GR cell line. HCD and EThcD were compared for **(A)** the resulting spectral score (Hyperscore) and **(B)** ID rates for the GR immunopeptidome. **(C)** Identified predicted binders (NetMHCpan 4.1, <1% binding rank) were analyzed for the percentage (%) of binders to the HLA-A, -B, and –C-alleles. Plotted are the cumulative percentages of predicted binders to both of the A and B alleles. N.B. Identified predicted binders can be predicted to bind multiple HLA-alleles. **(D)** Venn diagrams of the overlap between HCD- and EThcD-identified unique peptides for HLA-A*01:01, -A*03:01, –B*07:02 and –B27*05. **(E)** Sequence motifs of over-represented amino acids in identified peptides. Motifs on the top are enriched in EThcD-preferred peptides, whereas motifs on the bottom are less identifiable in the EThcD-preferred peptides compared to sequences identified by HCD alone + those identified by both HCD and EThcD (i.e. ‘HCD-preferred’). Motifs were extracted with Two-Sample Logo (p<0.05) for HLA-A*02:01 and –B*07:02 predicted binders (<1% binding rank). The height of each symbol is proportional to the difference in relative frequencies. **(F)** Percentage (%) of sequence coverage of HCD (b/y ion series) and EThcD (b/y, c/z, or the combination of ion series) for HLA-A*01:01, -A*03:01, –B*07:02 and –B*27:05 predicted binders (<1% binding rank).

Also, for the GR immunopeptidome a proportion of binders were preferentially identified by EThcD. The number of unique peptides uncovered solely by EThcD differed substantially per HLA allele: 417 (15%) for -A*03:01, 359 (20%) for -A*01:01, 1556 (30%) for -B*27:05, and 1360 (33%) for - B*07:02 (**Fig. 5D**), probably related to the presence of Arg within the peptide sequences (**Supplemental Fig. S2**). Consistent with the JY results, EThcD-preferred peptides for GR immunopeptides are also rich for Arg in the middle of the sequence for all alleles, except -A*01:01, which similar to -A*02:01 complex, does not allow a high representation of basic residues within its sequence (**Fig. 5E**). Across all alleles, EThcD fragmentation, when compared to HCD, demonstrated an increase of median sequence coverage due to the additional c/z ion series of: +15% for -A*03:01 and -B*07:02 and +20% for -A*01:01 and -B*27:05 (**Fig. 5F**). Among alleles there was a similar contribution of b/y and c/z ions to the sequence coverage, with the highest c/z contribution observed for -B*27:05, potentially due to its higher charge state distribution including more *z*=+3 ions (**Fig. 3B**). Of secondary interest, the number of unique peptides identified solely by HCD was found to be always below 9%, except for -A*03:01, with the number and percentages being per HLA allele: 361 (13%) for -A*03:01, 101 (6%) for -A*01:01, 410 (8%) for -B*27:05, and 366 (7%) for -B*07:02 (**Fig. 5D**). Here again, peptides with internal Pro residues and/or C-terminal Arg/Lys are preferentially identified by HCD, as are peptides harboring an internal Aspartic Acid in -A*01:01 peptides. All these features can be explained as Aspartic Acid potentially reducing the local charge of the peptides, and the Pro effect is well-known (68).

Finally, we included also an alike analysis on the HLA ligandomes of Jurkat cells. In line with the previous two cell lines, EThcD fragmentation boosted the confidence scoring by a percentage of 23% (**Fig. 6A**) and ID rate percentages by a total of 9.3% (**Fig. 6B**). Again no substantial differences were observed between HCD and EThcD in terms of allele-assignment of the identified peptides, with very similar observations for the well-known anchor residues (**Fig. 6C**). Next, the sequence features of the predicted binders below the 1% cut-off were compared between HCD and EThcD runs. For all alleles, a large number and proportion of EThcD-preferred peptides were identified: 395 (15%) for - A*03:01, 699 (36%) for -B*07:02 and 590 (33%) for -B*35:03 (**Fig. 6D**). Motif deconvolution of all EThcD-preferred peptides revealed again of strong presence of Arg residues in the middle of the peptide sequences (**Fig. 6E**). Consistent with the previous data, the median sequence coverage of the Jurkat immunopeptidome increased by 12.5-20% across peptides mapped to different HLA alleles by benefitting of the dual b/y and c/z ion series (**Fig. 6F**). Based on the consistent increase of 12.5-20% in sequence coverage by EThcD, our data indicate that EThcD has a substantial complementary value for immunopeptidome identification across a broad range of HLA types and may help to uncover important peptides missed by using solely HCD fragmentation.

**Figure 6.**
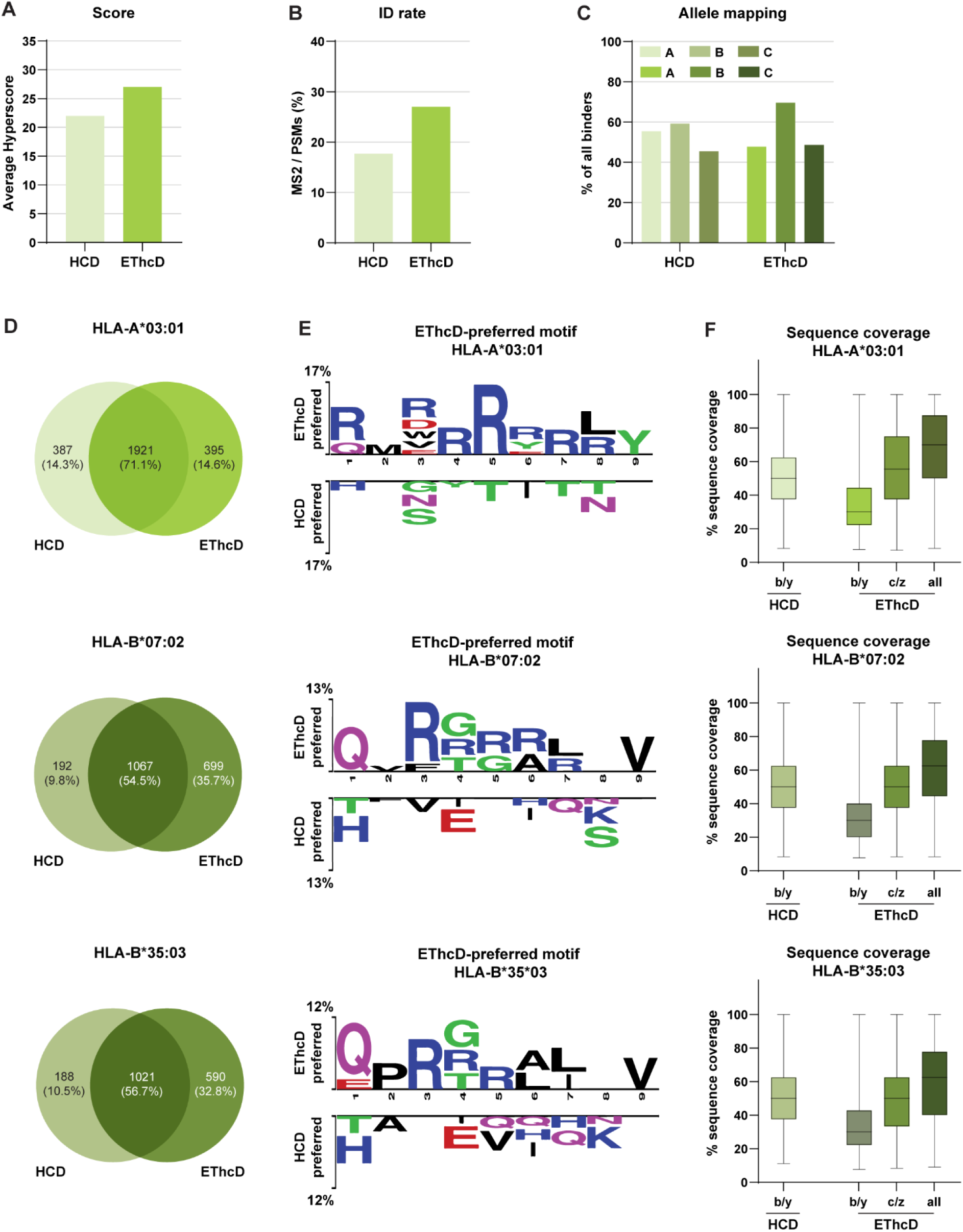
EThcD fragmentation extends the detectable immunopeptidome of the Jurkat cell line. HCD and EThcD were compared for **(A)** the resulting spectral score (Hyperscore) and **(B)** ID rates for the Jurkat immunopeptidome. **(C)** Identified predicted binders (NetMHCpan 4.1, <1% binding rank) were analyzed for the percentage (%) of binders to the HLA-A, -B, and –C-alleles. Plotted are the cumulative percentages of predicted binders to both of the B alleles. N.B. Identified predicted binders can be predicted to bind multiple HLA-alleles. **(D)** Venn diagrams of the overlap between HCD- and EThcD-identified unique peptides for HLA-A*03:01, –B*07:02 and –B35*03. **(E)** Sequence motifs of over-represented amino acids in identified peptides. Motifs on the top are enriched in EThcD-preferred peptides, whereas motifs on the bottom are less identifiable in the EThcD-preferred peptides compared to sequences identified by HCD alone + those identified by both HCD and EThcD (i.e. ‘HCD-preferred’). Motifs were extracted with Two-Sample Logo (p<0.05) for HLA-A*02:01 and –B*07:02 predicted binders (<1% binding rank). The height of each symbol is proportional to the difference in relative frequencies. **(F)** Percentage (%) of sequence coverage of HCD (b/y ion series) and EThcD (b/y, c/z, or the combination of ion series) for HLA-A*03:01, –B*07:02 and –B*35:03 predicted binders (<1% binding rank).

### EThcD expands the detectability of mono-methylated/di-methylated immunopeptides

Above, we demonstrated that EThcD enhances the identification of Arg-rich peptides across multiple HLA-alleles. EThcD has concomitantly been shown to have an inherent advantage when analyzing post-translational modifications (PTMs) (including site-localization) as the dual ion series provide richer, more confidently identified MS2 spectra, even in the presence of PTMs (35). To further investigate the potential of the new instrument, we explored whether EThcD could be used to detect HLA peptides harboring PTMs, and we focused here on Arg mono-methylation and di-methylation (**Supplemental Table 2**). Arginine methylation is a widespread protein PTM that plays crucial roles in a variety of fundamental cellular processes, such as gene transcription, epigenetics, RNA processing and genome maintenance (70, 71).

To this end, we performed PTM analysis to identify all mono-methylated/di-methylated peptides across both HCD and EThcD data from the JY and GR immunopeptidomes. Overall, we detected over hundred HLA peptides harboring Arg mono-methylation or di-methylation. In line with previous findings (37), almost all identified sequences harboring Arg methylation were HLA-B*07:02 binders (102/116, 88%) and harbored the PTM on P3 of the peptide sequence (**Fig. 7A-B**). From our JY and GR analysis, EThcD favored the identification of -B*07:02 binders, with high enrichment of Arg in P3 (**Fig. 4F and 5E**). As we hypothesized, a higher number of PTM sites were identified by EThcD compared to HCD for both the JY (+25%) and GR (+32%) immunopeptidome (**Fig. 7C**), corroborating that EThcD is important in analyzing peptide PTMs.

**Figure 7.**
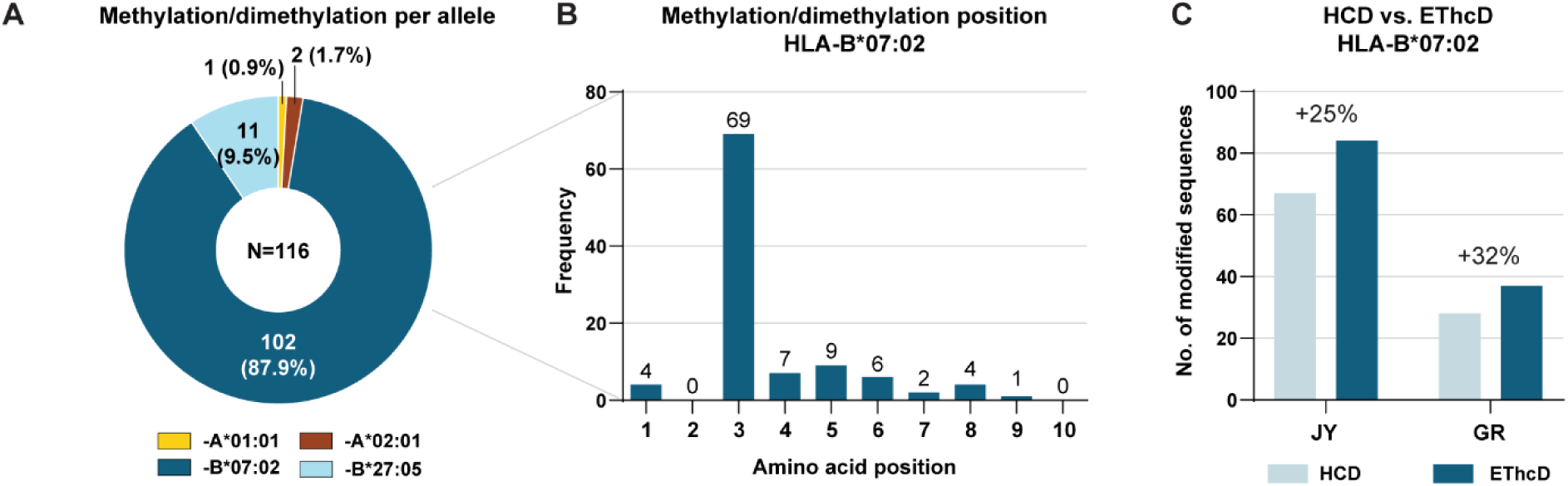
EThcD improves confidence in identification of post-translationally modified immunopeptides as exemplified by looking for Arg mono-methylated/di-methylated peptides in JY and GR samples. **(A)** The possible occurrence of mono- and di-methylation on Arg residues were searched for by using FragPipe in JY and GR samples. Their highly skewed distribution across predicted HLAs is depicted. **(B)** The positional frequency of mono- and di-methylations on Arg in HLA-B*07:02 predicted binders is strongly centered around P3. **(C)** Number of mono- and di-methylations on Arg residues in GR and JY for HLA-B*07:02 predicted binders were calculated as present in HCD (light blue bars) and EThcD (dark blue bars) data.

## Discussion

The identification of HLA-I-bound peptides by LC-MS/MS analysis has contributed significantly to our understanding of the antigen processing and presentation pathways mediating immune recognition of disease, and successfully led to the identification of targets for novel immunotherapeutic strategies including cancer vaccines (1, 13, 72). The selection of such actionable targets, however, is highly dependent on an extensive identification depth and accuracy of sequence assignment of the entire immunopeptidome. The study presented here had three aims. First, to explore the potential of the new Orbitrap Excedion Pro mass spectrometer and its ability to perform ETD/EThcD for the analysis of non-tryptic HLA-I and elastase-digested peptides, exploiting the fast ETD reaction within the IRM (**Fig. 1**). Second, to demonstrate the inability of collision-based fragmentation methods, currently recognized as the golden standard for fragmentation, to identify a proportion of peptides due to HLA allele-specific biases in immunopeptide. Third, demonstrate that EThcD enables complementarity in identified sequences and superiority in sequence coverage across HLAs, allowing higher sequence assignment accuracy, compared to CID/HCD. We first optimized EThcD method parameters using an elastase-digested HeLa cell lysate as an immunopeptidome surrogate (**Supplemental Table 1** and **Fig. 2**), which we then used to analyze immunopeptidomes derived from several of the globally most frequent HLA alleles (i.e. HLA-A*01:01, -A*02:01, A*03:01, and -B*07:02) (63–65) representing a wide range of charge states, amino acid distributions and biophysical properties (42).

Our re-analysis of multiple publicly allele-matched immunopeptidome datasets acquired by HCD (**Supplemental Table 3 and Fig. 3**; (14, 50, 53, 54)) hinted to HCD limitations especially for HLA-A*02:01 peptides, an HLA allele expressed in a large part of the world population and at the center of immunotherapy development (73), (**Fig. 3D-E**) leading to potential identification loss and inaccuracy of assignment.

We then evaluated the fine-tuned EThcD methods to analyze immunopeptidomes of JY, GR and Jurkat cell lines (**Supplemental Table 2**). For the JY cell line, we tested the effect of excluding (M1 method) or including (M2 method) z=+1 precursors and different scan *m/z* ranges. We found that, in this mass analyzer, unlike other platforms (52), exclusion of *z*=+1 precursors provides higher benefits, which can be explained by the narrower, sensitive scan range and a higher MS2 efficiency due to the exclusion of *z*=+1 potential contaminants.

We consistently found higher sequence coverages across HLA-bound immunopeptidomes by EThcD compared to HCD (**Figs 4G, 5F, 6F**), which is the likely explanation for a high proportion of translatable MS2 spectra across all cell lines (ID rate gain; ranging between +9.3% to +16.4%; **Fig 4B, 5B, 6B**), making up for any expected reduction in duty cycle of EThcD compared to CID/HCD workflows.

We next explored across all alleles whether the increased sequence coverage and identification score confidence by EThcD would translate into unique identifications that are consistently missed by HCD (i.e. ‘EThcD-preferred peptides’) (**Fig. 4F, 5E** and **6E**), being mostly hampered by Arg residues centered in the middle of the sequence, leading to a reduction in peptide backbone fragmentation (67). The exception in this finding were the HLA-A*02:01-bound peptides, which did not show high enrichment for Arg residues (**Fig. 4F**). We postulate that EThcD gain for HLA-A*02:01 could instead be driven by the baseline lower HCD fragmentation efficiency compared to other alleles as observed by the high average sequence gap of 4.5 (**Fig. 3E**), resulting in unassigned MS2 spectra. Of note, the HLA-A*02:01 motif displays a strong presence of Leu at P2 and the C-term (**Supplemental Fig. S2**). Leu is isobaric to isoleucine (Iso) and is not readily distinguishable by HCD, increasing the potential misassignment of these peptides by HCD. While we haven’t studied it in this manuscript, EThcD can aid distinguishing between Leu/Iso by generation of secondary ion series (i.e. W) in HLA-A*02:01 peptides (74).

We further strengthened our hypothesis that a substantial portion of immunopeptides cannot yield sufficient diagnostic fragments when using HCD as the sole fragmentation scheme by comparing EThcD-preferred peptides from our JY data to a much larger JY dataset (>19000 unique peptides) acquired in different laboratories (**Supplemental Table 3**). We noted that a meaningful portion (31% and 39% for HLA-A*02:01 and –B*07:02, respectively) of the identified EThcD-preferred peptides were not detectable in this large JY HCD data.

Lastly, the benefit of EThcD for PTM identification and localization have been demonstrated before for immunopeptidomics (35). In present study, we wanted to combine our findings in EThcD superiority in identifying Arg-rich sequences to the study of mono- and di-methylation of Arg (**Supplemental Table 2**). We showcased that most identified PTMs were mapped back to HLA-B*07:02 (**Fig. 7A**), have an allele-specific sub-anchor preference of Arg in P3 (**Fig. 7B**), and that EThcD expectedly enhances the identification between 25-32% of these PTMs sites compared to HCD (**Fig. 7C**).

## Conclusion

Here, we describe the first applications on a new mass spectrometer termed the Orbitrap Excedion Pro. This hybrid benchtop Orbitrap-based instrument has enabled a novel approach to perform ETD, HCD, and hybrid fragmentation, i.e., EThcD. By performing the ETD reaction in the ion routing multipole (IRM) it results in a fast and efficient electron-induced fragmentation capabilities, thus reducing the duty cycle utilization gap when compared to HCD alone. While this is a significant step forward for non-tryptic, lower charged peptides, we envision that further technological steps would be beneficial to efficiently couple EThcD to faster scanning, more sensitive analyzers (52, 75, 76). We demonstrate the performance of this instrument, in particular for the analysis of a wide range immunopeptidomes and the non-tryptic elastase digest. EThcD holds imperative value for the field of immunopeptidomics demonstrated by the higher sequence coverages (thus reducing sequence gaps), increased ID rates and more confident scores across a wide range of immunopeptides with variable physicochemical properties leading to systematically uncovering a set of immunopeptides consistently missed with HCD. We thus postulated that no matter what the speed or sensitivity of the analyzer, when using HCD/CID fragmentation alone, a subset of immunopeptides will remain undiscovered. Furthermore, relying solely on CID/HCD for therapeutic target discovery warrants caution, as lower sequence coverages hinder accuracy of sequence assignment by both database as well as *de novo* searches. This may lead to loss of identification or extensive validation of clinically relevant antigens; especially as shown for the HLA-A*02:01-associated peptidome. We envision that, upon wide implementation in the field of antigen-based immunotherapies, EThcD will enable deeper, complementary and more accurate identification of clinically relevant antigens, including non-canonical targets (77) for pioneering immunotherapeutic strategies. Furthermore, another important gain will be observed for other hard to fragment PTMs and the localization of PTMs, including glycosylation and phosphorylation (among others). Beyond HLA class I, HLA class II peptides (78) and other, non-tryptic longer peptides in the field of middle-down proteomics (79–81) and antibody *de novo* sequencing (31, 82) will benefit from the above discussed instrument advancements, as well as other applications where non-tryptic peptides (e.g. the field of *de novo* sequencing) are utilized.

## Data availability

The mass spectrometry immunopeptidomics data generated for this study have been deposited to the ProteomeXchange Consortium (http://proteomecentral.proteomexchange.org) via the PRIDE partner repository (83) with the dataset identifier PXDxxxxxx.

## Supplemental data

This article contains supplemental data (figures and tables).

## Author contributions

*Conceptualization*: A.J.R.H., A.L.K., F.M., H.K., K.L.F.; *Methodology*: A.L.K., F.M., K.L.F., P.K.; *Software*: J.H., K.L.F., P.K.; *Validation*: A.L.K., F.M., H.C.R., K.L.F., P.K.; *Formal analysis*: A.L.K., F.M., H.C.R., K.L.F.; *Investigation*: A.L.K., F.M., H.C.R., P.K.; *Resources*: A.L.K., C.W., F.M., H.C.R., H.K., J.H., K.L.F., P.K.; *Data curation*: A.L.K., F.M., K.L.F.; *Writing – original draft*: A.J.R.H., A.L.K., F.M.; *Writing – Review & Editing*: A.J.R.H., A.L.K., C.W., F.M., H.C.R., H.K., K.L.F., P.K.; *Visualization*: A.L.K., F.M.; *Supervision*: A.J.R.H., F.M., K.L.F.; *Project administration*: A.J.R.H., A.L.K., F.M., H.K., K.L.F.; *Funding acquisition*: A.J.R.H..

## Abbreviations

AGC: Automatic gain control
*z*: Charge
CID: Collision induced dissociation
ETD: Electron-transfer dissociation
EThcD: Electron-transfer/higher-energy collision dissociation
HCD: Higher-energy collision dissociation
HLA-I: Human leukocyte antigen class I
IRM: Ion routing multipole
LC-MS/MS: Liquid chromatography coupled to tandem mass spectrometry
*m/z*: Mass over charge
NCE: Normalized collision energy
PSMs: Peptide spectrum matches
RF: Radio frequency
RP: Reversed phase
SA: Supplemental activation

