## Supplemental Figure 1, 2 and 3 for "Increased EThcD efficiency on the hybrid Orbitrap Excedion Pro Mass Analyzer extends the depth in identification and sequence coverage of HLA class I immunopeptidomes"

**Table of contents**

- Supplemental Fig. S1
- Supplemental Fig. S2
- Supplemental Fig. S3
- References

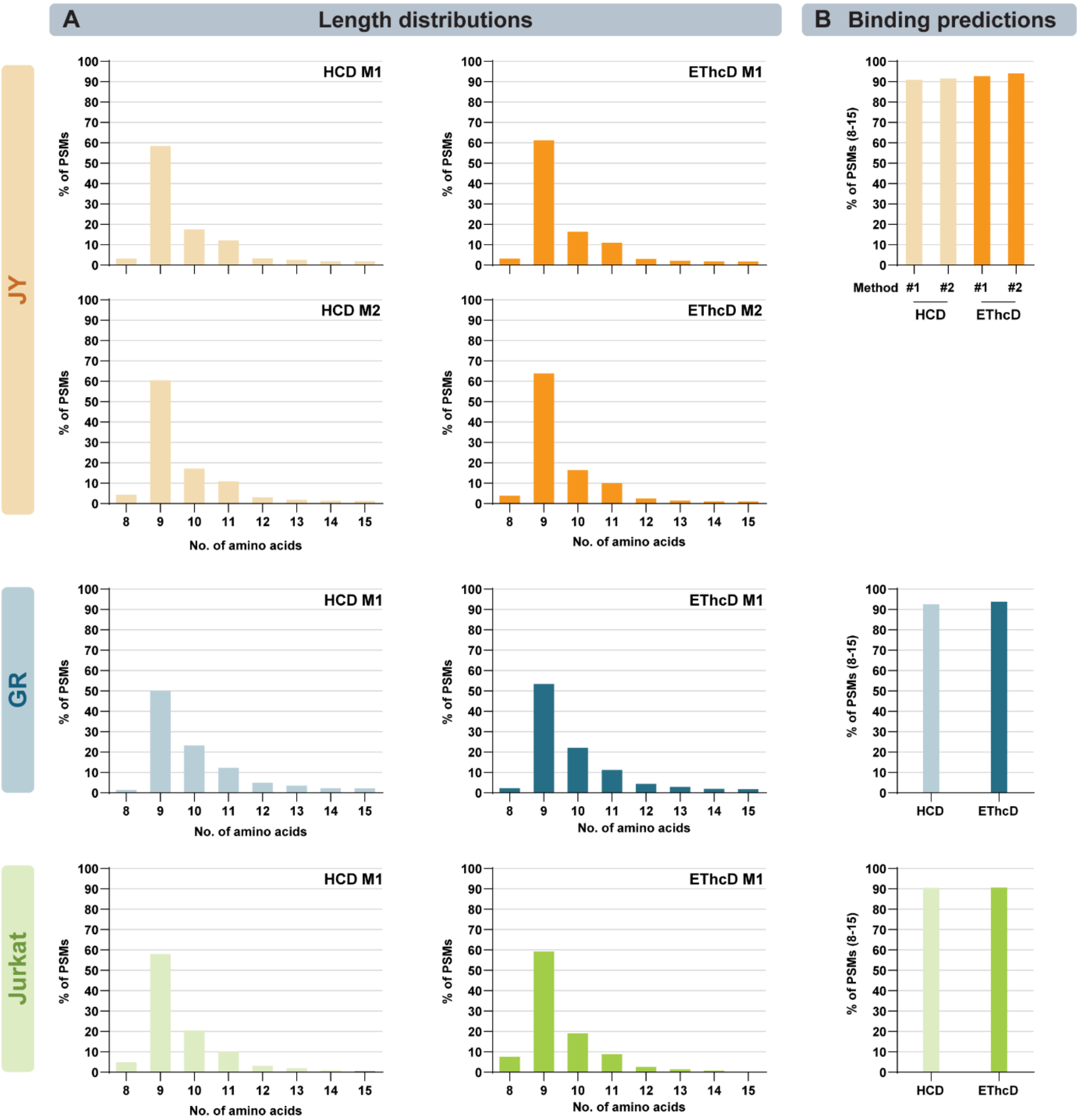


**Supplemental Figure S1 | Generic features of the detected immunopeptidomes. Peptide length distribution and HLA binding prediction analyses indicate high quality enrichment of HLA-bound peptides. (A)** Length distribution analysis of all identified PSMs (8-15-mers) in HLA peptide-enriched samples of JY (orange), GR (blue) and Jurkat (green) cells. Each bar represents the percentage of all PSMs that has a certain length (in amino acids). At the top right the used MS2 method is shown. **(B)** Percentage (%) of predicted HLA binders in the dataset of all identified PSMs (8-15-mers) in each sample by NetMHCpan 4.1 (<2% binding rank).

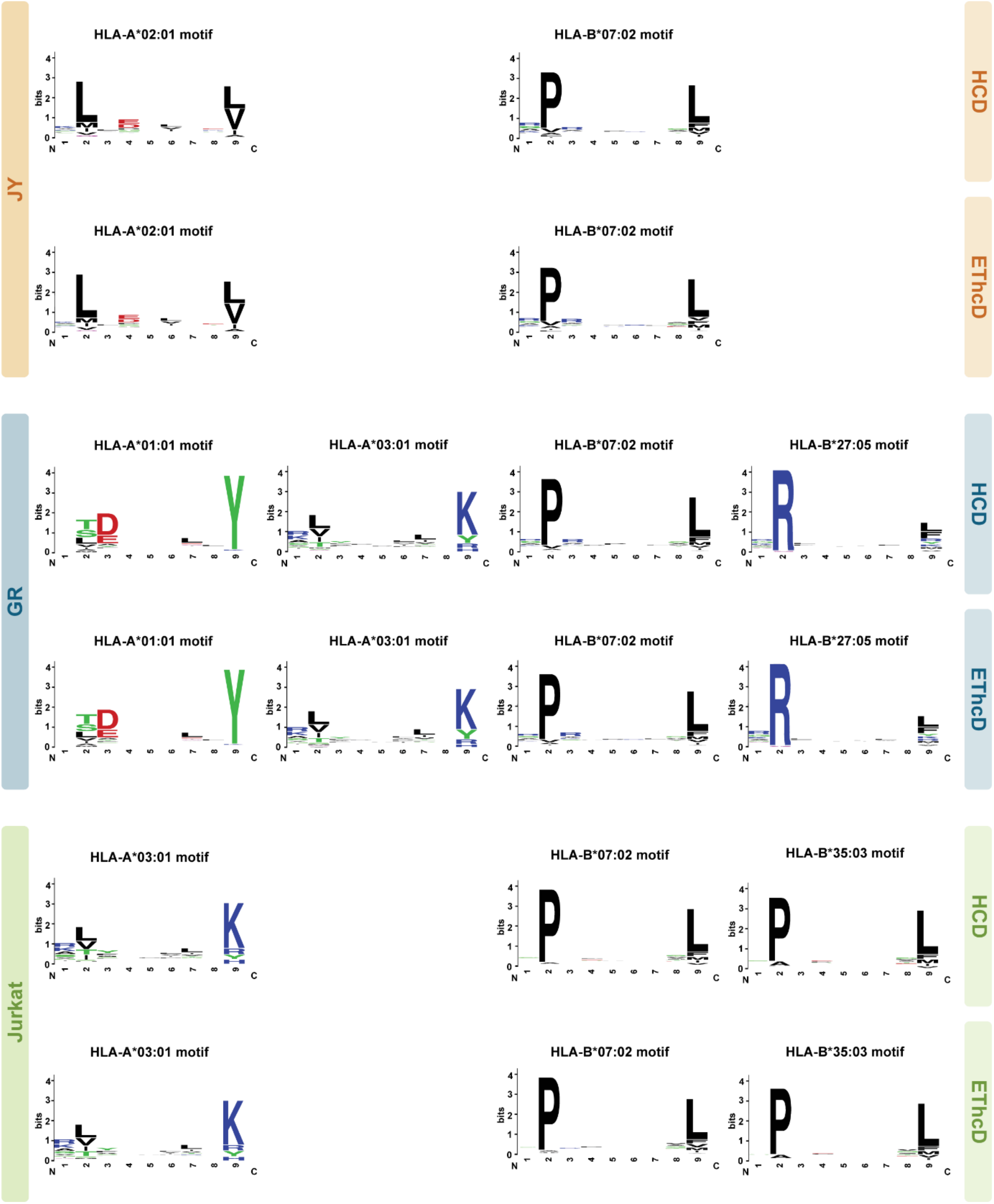


**Supplemental Figure S2 | Sequence motif analysis of all identified predicted HLA binders in JY, GR and Jurkat immunopeptidomes.** Sequence motifs were extracted from all predicted binders (NetMHCpan 4.1, <1% binding rank) with WebLogo. The height of each symbol indicates the relative frequency of each amino acid in the dataset at that given position. All motifs are highly alike those reported earlier for each of the allele investigated here (1).

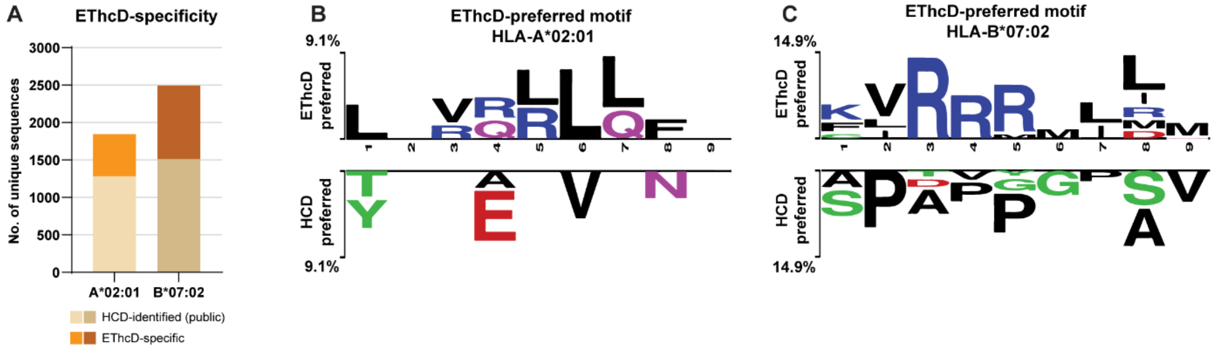


**Supplemental Figure S3 | Validation of the observed preference of EThcD-specific identifiable immunopeptides by cross-referencing with an extended large dataset of publicly available HCD immunopeptidomics data. (A)** EThcD-preferred peptides predicted to bind HLA-A*02:01 or –B*07:02 were extracted and cross-referenced to a large dataset of JY HCD immunopeptidomics data (**Supplemental Table 3**). The total bar height corresponds to the total number of EThcD-preferred peptides identified in our data. The brown-toned 'HCD-identified (public)' bars correspond to the number of EThcD-preferred peptides that were also identified in the public HCD dataset. The orange-toned 'EThcD-specific' bars indicate the number of peptides that were not earlier detected by using HCD. The EThcD-specific peptides binding to **(B)** HLA-A*02:01 and **(C)** HLA-B*07:02 were extracted, and sequence logos were generated with TwoSampleLogo ($p<0.05$). Displayed are sequence motifs of over-represented amino acids in EThcD-specific nonamers. Motifs on the top are enriched in EThcD-preferred peptides, whereas motifs on the bottom are less identifiable in the EThcD-preferred peptides compared to sequences identified by HCD alone + those identified by both HCD and EThcD (i.e. 'HCD-preferred'). The height of each symbol is proportional to the difference of relative frequencies.
